# SARS-CoV-2 Mac1 is required for IFN antagonism and efficient virus replication in mice

**DOI:** 10.1101/2023.04.06.535927

**Authors:** Yousef M. Alhammad, Srivatsan Parthasarathy, Roshan Ghimire, Joseph J. O’Connor, Catherine M. Kerr, Jessica J. Pfannenstiel, Debarati Chanda, Caden A. Miller, Robert L. Unckless, Sonia Zuniga, Luis Enjuanes, Sunil More, Rudragouda Channappanavar, Anthony R. Fehr

**Affiliations:** Department of Molecular Biosciences, University of Kansas, Lawrence, Kansas 66047, USA; Department of Veterinary Pathobiology, Oklahoma State University, Stillwater, Oklahoma, United States; National Center of Biotechnology (CNB-CSIC), Campus Universidad Autónoma de Madrid, Madrid, Spain

**Author notes:** These authors contributed equally to this work (Yousef M. Alhammad, Srivatsan Parthasarathy, Roshan Ghimire).

**Keywords:** Coronavirus, SARS-CoV-2, Non-Structural Proteins, Macrodomain, ADP-ribosylation, Interferon, Cytokines, Inflammatory Monocytes, Innate Immunity

## Abstract

Several coronavirus (CoV) encoded proteins are being evaluated as targets for antiviral therapies for COVID-19. Included in this set of proteins is the conserved macrodomain, or Mac1, an ADP-ribosylhydrolase and ADP-ribose binding protein. Utilizing point mutant recombinant viruses, Mac1 was shown to be critical for both murine hepatitis virus (MHV) and severe acute respiratory syndrome (SARS)-CoV virulence. However, as a potential drug target, it is imperative to understand how a complete Mac1 deletion impacts the replication and pathogenesis of different CoVs. To this end, we created recombinant bacterial artificial chromosomes (BACs) containing complete Mac1 deletions (ΔMac1) in MHV, MERS-CoV, and SARS-CoV-2. While we were unable to recover infectious virus from MHV or MERS-CoV ΔMac1 BACs, SARS-CoV-2 ΔMac1 was readily recovered from BAC transfection, indicating a stark difference in the requirement for Mac1 between different CoVs. Furthermore, SARS-CoV-2 ΔMac1 replicated at or near wild-type levels in multiple cell lines susceptible to infection. However, in a mouse model of severe infection, ΔMac1 was quickly cleared causing minimal pathology without any morbidity. ΔMac1 SARS-CoV-2 induced increased levels of interferon (IFN) and interferon-stimulated gene (ISG) expression in cell culture and mice, indicating that Mac1 blocks IFN responses which may contribute to its attenuation. ΔMac1 infection also led to a stark reduction in inflammatory monocytes and neutrophils. These results demonstrate that Mac1 only minimally impacts SARS-CoV-2 replication, unlike MHV and MERS-CoV, but is required for SARS-CoV-2 pathogenesis and is a unique antiviral drug target.

**SIGNIFICANCE:** All CoVs, including SARS-CoV-2, encode for a conserved macrodomain (Mac1) that counters host ADP-ribosylation. Prior studies with SARS-CoV-1 and MHV found that Mac1 blocks IFN production and promotes CoV pathogenesis, which has prompted the development of SARS-CoV-2 Mac1 inhibitors. However, development of these compounds into antivirals requires that we understand how SARS-CoV-2 lacking Mac1 replicates and causes disease *in vitro* and *in vivo*. Here we found that SARS-CoV-2 containing a complete Mac1 deletion replicates normally in cell culture but induces an elevated IFN response, has reduced viral loads *in vivo*, and does not cause significant disease in mice. These results will provide a roadmap for testing Mac1 inhibitors, help identify Mac1 functions, and open additional avenues for coronavirus therapies.

## INTRODUCTION

Coronaviruses (CoVs) belong to the family *coronaviridae* and possess a large, positive-sense RNA genome. The subfamily *coronavirinae* is further subdivided into α, β, γ and δ-CoVs, though only the α and β-CoVs include viruses that infect humans. Prior to the 21^st^ century CoVs were predominantly known to cause mild respiratory disease in humans (1). However, with the emergence of SARS-CoV, MERS-CoV, and most recently SARS-CoV-2, it is now well-established that CoVs are implicated in severe human respiratory conditions and are a serious threat to human health.

*<u>Co</u>*rona*v*irus *i*nfectious *<u>d</u>*isease (COVID-19) caused by SARS-CoV-2 is responsible for the pandemic that has resulted in over 6 million deaths worldwide (WHO). In cases of severe COVID-19, SARS-CoV-2 induces a robust pro-inflammatory cytokine response, or cytokine storm, in the host leading to the development of acute respiratory distress syndrome (ARDS) and in some cases multiple organ pathologies (2). Introduction of SARS-CoV-2 mRNA vaccines have drastically increased antiviral immunity and has reduced the fatality caused by SARS-CoV-2 (CDC). However, many elderly or immunocompromised people have ineffective responses to vaccines (3), and with the rate of emergence of new SARS-CoV-2 variants like Omicron (BA.2, BA.4 and BA.5) there is an urgent need to identify novel antiviral drugs. Currently, a few antiviral drugs such as Veklury (Remdesivir) (4, 5) and Lagevrio (molnupiravir) (5) both of which target the CoV polymerase (nsp12); and Paxlovid (nirmatrelvir and ritonavir) (6), which targets the main protease (nsp5), have been utilized in hospitals to treat COVID-19 patients under adverse conditions. However, it remains important to identify novel drug targets to expand the pool of anti-CoV therapies that will be needed to account for drug-resistance, provide additional options for treatment, and better understand the replication processes of CoVs.

All CoVs encode a conserved set of 15-16 non-structural proteins that direct the formation of the replication transcription complex (RTC) and carryout the process of RNA transcription and replication, making these proteins important targets for antiviral therapies. While much progress has been made in identifying the functions of many of the non-structural proteins, we still lack a complete understanding of how these proteins contribute to RNA replication and evasion of the host immune response. Non-structural protein 3 (nsp3) is the largest non-structural protein encoded in the CoV genome and consists of several modular protein domains, such as the papain-like protease (PLP) domain. Included in these domains of nsp3 are 3 tandem macrodomains (Mac1, Mac2 and Mac3). Mac1 is conserved throughout all CoVs unlike Mac2 and Mac3 (7-12). Notably, homologs of Mac1 are also found in other viruses like alphaviruses, hepatitis E virus, and rubella virus, suggesting it could play an important role in the replication of a subset of positive-strand RNA viruses (13, 14). Structurally, macrodomains are characterized by the presence of a conserved three-layered α/β/α fold. Biochemically, the conserved viral macrodomain binds to ADP-ribose moieties with high affinity (15, 16) and in some cases can hydrolyze the bond between ADP-ribose and proteins, reversing ADP-ribosylation, a common post-translational modification (15, 17-20).

ADP-ribosylation is catalyzed by ADP-ribosyltransferases (ARTs/PARPs) using NAD^+^ as the substrate (21). ADP-ribose subunits can be added to proteins as single subunits in a process termed mono-ADP-ribosylation (or MARylation) or as a polymer of ADP-ribose subunits forming a chain in a process termed poly-ADP-ribosylation (or PARylation). Notably, several of the MARylating PARPs are interferon stimulated genes (ISGs) and demonstrate antibacterial and antiviral properties (22-25). These results highlight the importance of ADP-ribosylation as a putative antiviral host response and viral macrodomains as an evolutionary adaptation by certain viruses to counter this host response (17, 19, 20). Therefore, it is of interest to better understand how viral macrodomains counter PARP activity and contribute towards viral infection and pathogenesis.

The recent body of research has identified Mac1 as a viral factor necessary for CoV replication and pathogenesis in multiple animal models of infection (18, 26). Most of these studies have utilized a point mutant of Mac1 where a conserved asparagine residue (N1347 in MHV-JHM) was mutated to an alanine. This mutation dramatically reduces the ability of Mac1 to hydrolyze MAR from target proteins (16, 18, 27-29). The SARS-CoV-1 Mac1 asparagine-to-alanine mutant virus (N1040A) had minimal to no impact on replication in transformed cells but was sensitive to IFN pre-treatment and induced robust IFN and pro-inflammatory cytokine production and caused minimal disease in mouse models of infection (18, 30). Similar results were seen with the corresponding MHV Mac1 mutant virus (N1347A), though growth defects have been observed in some cell types with these viruses (31-33). Importantly MHV-N1347A replication increased upon PARP inhibition or knockdown of PARP12 or 14, while WT virus was unaffected.

Similarly, IFN induction following infection with MHV N1347A was nearly eliminated upon PARP inhibition and in PARP14 knockout cells. These results demonstrate that Mac1 function countered the action of PARP mediated ADP-ribosylation (32). Apart from N1347A, we found that another unique mutation, D1329A, a residue which is critical for the ADP-binding activity of macrodomains, replicated poorly in multiple cell types. Additionally, we were unable to recover an MHV double mutant virus, D1329A/N1347A, indicating that Mac1 may be critical for CoV replication. These results demonstrate that Mac1 has multiple functions that can promote viral replication and block host interferon responses (33).

These combined studies have prompted several groups to begin screening for and developing SARS-CoV-2 Mac1 inhibitors that could potentially be used therapeutically to treat patients infected with SARS-CoV-2 or other emerging CoVs (34-41). However, before testing any of these inhibitors for their antiviral activity, it is imperative to determine the role and functions of Mac1 in SARS-CoV-2 replication, pathogenesis, and the host immune response. Here we have created a complete SARS-CoV-2 Mac1 deletion virus and characterized its replication and immune modulating properties. These results provide new insights into SARS-CoV-2 biology, the innate immune response to infection, and will provide a roadmap for future testing of Mac1 inhibitors for antiviral activity.

## RESULTS

### SARS-CoV-2 Mac1 deletion virus infectious virus was easily recovered while Mac1 deletion viruses in other *β*-CoVs were not recovered

We recently identified several Mac1 mutations in murine hepatitis virus strain JHM (MHV-JHM) that were unrecoverable from a bacterial artificial chromosome (BAC) based reverse genetic system (33). These results indicated that Mac1 may be critical for MHV replication. As point mutations could result in toxic unfolded proteins, we created an MHV-JHM Mac1 deletion BAC to confirm our prior results. As expected, we were unable to recover infectious virus (Fig. S1A-B) from the Mac1 deletion BAC, further indicating that Mac1 is critical for MHV-JHM replication. We next created a complete deletion of Mac1 in MERS-CoV, and again we were unable to recover infectious virus, indicating that Mac1 is also critical for MERS-CoV replication (Fig. S1A-B). We hypothesized that Mac1 might be essential for the replication of all CoVs, so we engineered a Mac1 deletion (ΔMac1) into a SARS-CoV-2 BAC (Wuhan strain) to provide additional evidence for our hypothesis. However, unlike MHV-JHM or MERS-CoV, this virus was easily recoverable (Fig. S1A-B). This result indicates that there are stark differences in the requirement for Mac1 between SARS-CoV-2 and other *β*-CoVs.

### SARS-CoV-2 ΔMac1 replicates like WT virus in most cell types

Next, we assessed the ability of SARS-CoV-2 ΔMac1 to replicate in several cell types susceptible to SARS-CoV-2. In Vero E6 cells ΔMac1 replicated like WT virus cells at both low (Fig. 1A) and high (Fig. 1B) multiplicity of infection (MOI), indicating that Mac1 is not required for general virus replication. Vero E6 cells lack the ability to produce IFN-I, and MHV-JHM Mac1 mutant viruses are more attenuated in cells that maintain the ability to produce IFN-I (32). Thus, we hypothesized that SARS-CoV-2 ΔMac1 may be attenuated in either A549-ACE2 (alveolar epithelial cells) or Calu-3 cells (bronchial epithelial cells) that have a functional IFN system. ΔMac1 replicated equally to WT virus in A549-ACE2 cells (Fig. 1C), however there was a mild, ∼2-3-fold reduction in ΔMac1 titers in Calu-3 cells compared to WT virus at both low (Fig. 2A-B) and high MOI (Fig. 2C-D). We further observed only mild, if any, reduction in viral N protein when analyzed by immunoblotting and we observed roughly equal levels of both N protein and nsp3 staining by confocal microscopy (Fig. 2E-F) in Calu-3 cells infected by WT and ΔMac1. To evaluate the relative fitness of ΔMac1 compared to WT virus, we performed a competition experiment where we co-infected Calu-3 cells with WT and ΔMac1 at ratios of 1:1 and 1:9, respectively, and followed these viruses over the course of 4 passages. Virus was collected at approximately 36 hpi after each passage to isolate virus during active replication, and not after peak replication has been reached. To distinguish between WT and ΔMac1 viruses, we used semi-quantitative RT-PCR with primers set outside of Mac1 that produce different sized PCR products from each virus. First, using BAC DNA, we found that the ratio of these bands correlated with the ratio of input BAC (Fig. S2A-B), indicating that this method could faithfully define the relative abundance of each virus following passaging. We found that after 4 rounds of passaging ΔMac1 had not been outcompeted by WT virus as the ratios of these two viruses stayed relatively stable over the entire experiment (Fig. S2C-D), though WT virus was starting to increase in abundance in the 9:1 (ΔMac1:WT) sample at passage 4. In total, these results indicate that ΔMac1 generally replicates like WT virus but has a mild replication defect in Calu-3 cells, though it has similar fitness as WT virus in Calu-3 cells.

**Fig. 1.**
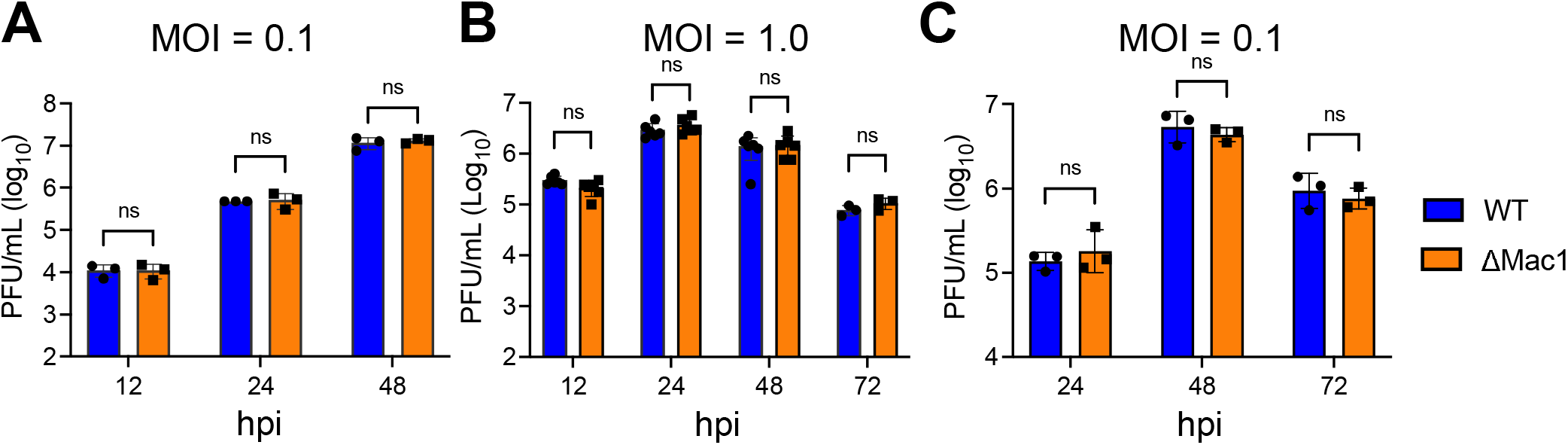
SARS-CoV-2 Mac1 deletion virus replicates normally in Vero E6 and A549-ACE2 cells. vroE6 (A-B) and A549-ACE2 (C) cells were infected with SARS-CoV-2 WT and ΔMac1 at an MOI of (A,C) and 1 (B) PFU/cell. Both cell-associated and cell-free virus was collected at indicated time nts and virus-titers were determined by plaque assays. Data shown is one experiment representative of three independent experiments. *n* = 3 per group for each experiment. ns – not significant.

**Fig. 2.**
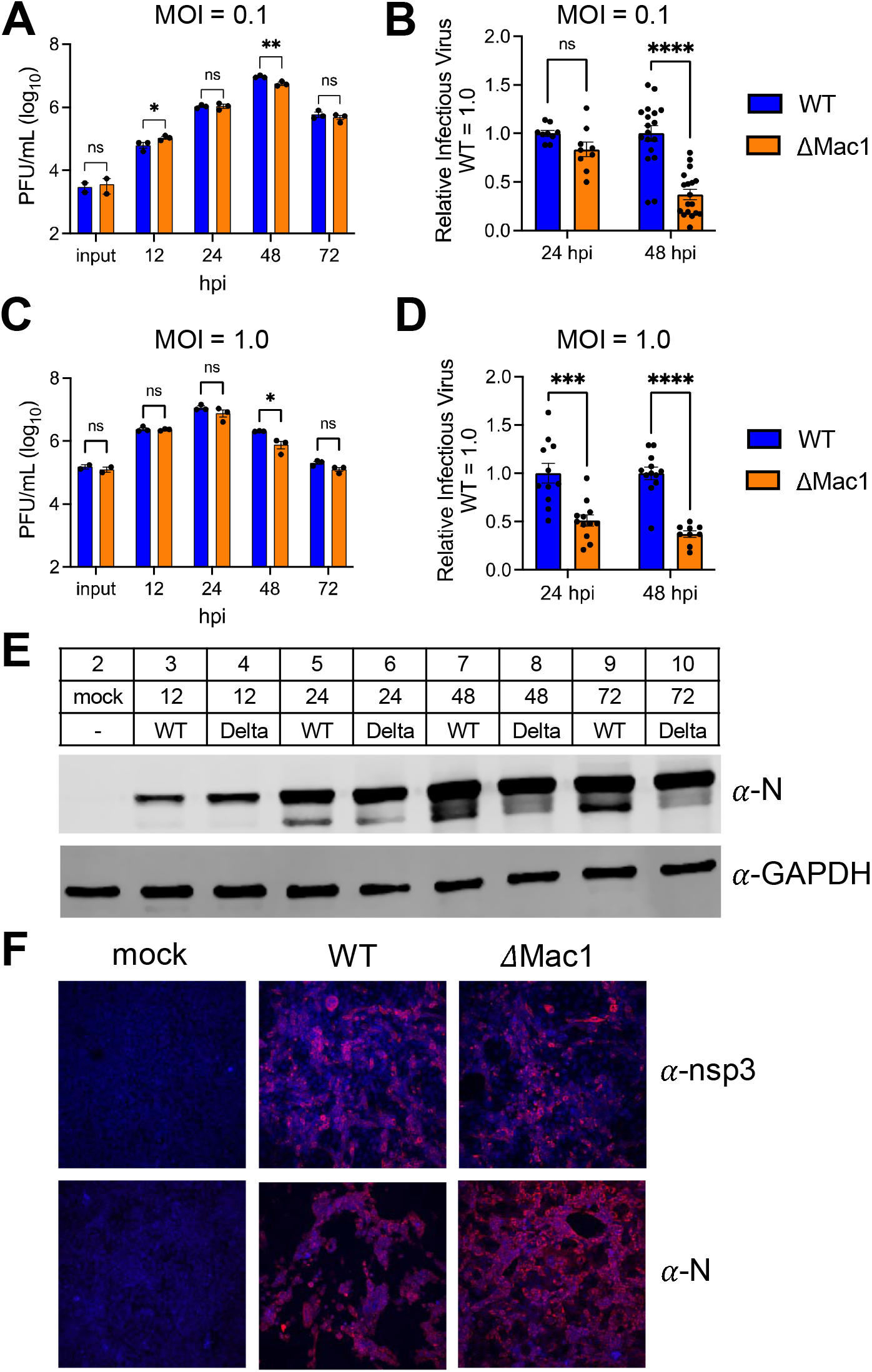
SARS-CoV-2 has a mild replication defect in Calu-3 cells. (A-D) Calu-3 cells were infected with -CoV-2 WT and ΔMac1 viruses at both low (A-B) and high (C-D) MOI. Both cell-associated and cell-irus was collected at indicated times and virus titers were determined by plaque assay. The data in A re from one experiment representative of at least 3 independent experiments. *n* = 3 per group. The s of all combined experiments where the average WT values from each experiment were normalized at 24 and 48 hpi are shown in B and D. Each point represents a separate biological replicate. (E-F) 3 cells were infected at an MOI of 1 PFU/cell as described above and cell lysates were collected, and rotein levels were determined by immunoblotting (E) or cells fixed at 24 hpi were co-stained with DAPI ither anti-nsp3 or anti-N, and then analyzed by confocal microscopy at 20X magnification (F). The data shows data from one representative experiment of two independent experiments.

### SARS-CoV-2 ΔMac1 is more sensitive to IFN-γ pre-treatment than WT virus

We previously found that IFN-/*β* pretreatment more effectively reduced MHV Mac1 mutant virus replication than WT virus replication in primary macrophages, likely due to the significant upregulation of PARP enzymes (32). Thus, we tested the ability of IFN-/*β* pre-treatment to impede SARS-CoV-2 WT and ΔMac1 infection in Calu-3 cells. We found that adding increasing amounts of IFN-/*β* to cells 18 hours before infection reduced ΔMac1 replication to the same level as WT virus in Calu-3 cells (Fig. 3A), indicating that SARS-CoV-2 ΔMac1 is not more sensitive to IFN-/*β* than WT virus. We hypothesized that SARS-CoV-2 may be too sensitive to IFN-/*β* to distinguish any difference in the replication of WT and ΔMac1 viruses. Therefore, we next used IFN-γ, which induces a smaller number of ISGs and has reduced antiviral activity against SARS-CoV-2 compared to IFN-/*β* (42), but still induces the expression of PARP enzymes (Fig. S3) (43). In contrast to results with IFN-/*β*, pre-treatment of cells with increasing concentrations of IFN-γ led to more robust inhibition of ΔMac1 than WT virus when cells were infected at an MOI of 0.1 and harvested at 48 hpi (Fig. 3B). The fold-difference in replication between WT and ΔMac1 ranged from 3-fold with no IFN-γ, which is consistent with results in Fig. 2, to ∼20-fold reduction in replication of ΔMac1 compared to WT virus when cells were pretreated with 500 units of IFN-γ (Fig. 3B). These results demonstrate that IFN-γ pre-treatment of Calu-3 cells more effectively impedes the replication of SARS-CoV-2 ΔMac1 compared to WT virus.

**Fig. 3.**
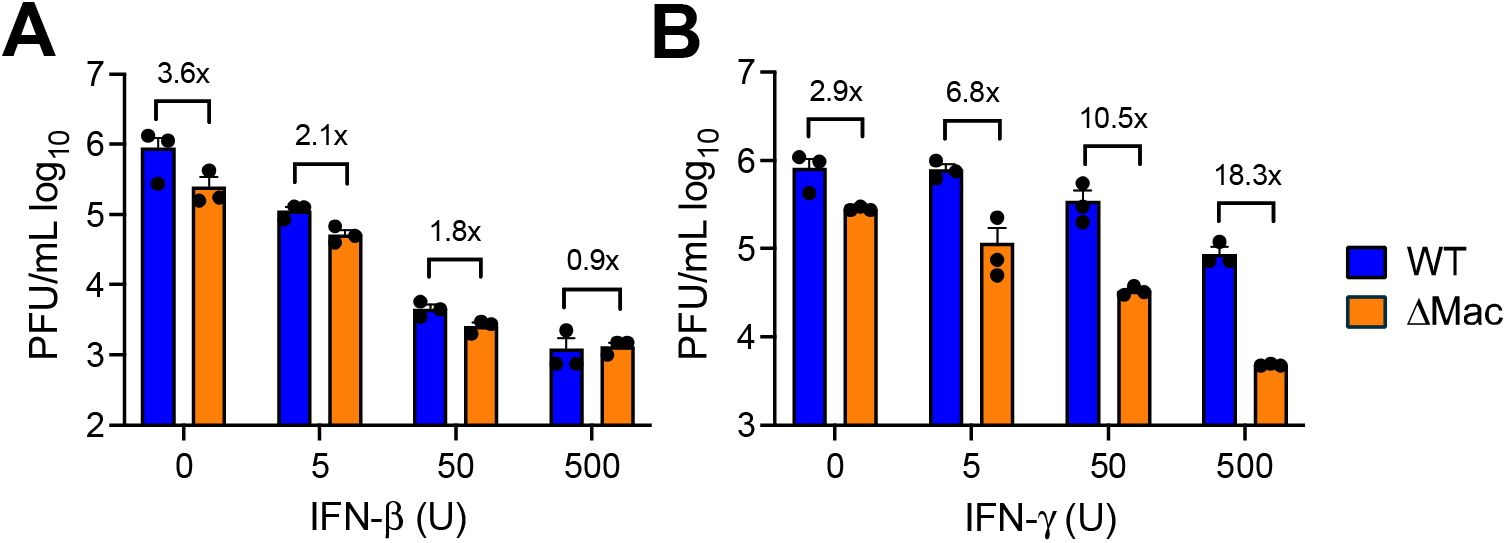
IFN-γ, but not IFN-β, pretreatment enhances replication defect of ΔMac1 in Calu-3 cells. Calu-3 cells were pretreated for 18 h with increasing concentrations (0, 5, 50, and 500 units) of IFN-β (**A**) and IFN-γ (**B**), then infected with either SARS-CoV-2 WT or ∆Mac1 at an MOI of 0.1 PFU/cell. Cells were collected at 48 hpi and titers were determined by plaque assay. Fold differences between WT and ΔMac1 are indicated at each amount of IFN. The data shown are of one experiment representative of two (A) and three (B) independent experiments. *n* = 3 for each group.

### SARS-CoV-2 ΔMac1 induces increased IFN and cytokine responses in cell culture

Next, we tested SARS-CoV-2 ΔMac1 for its ability to induce IFNs and pro-inflammatory cytokines in cell culture, as has previously been shown for Mac1 mutants in SARS-CoV and MHV (18, 32). We found that in both Calu-3 (Fig. 4A) and A549-ACE2 cells ΔMac1 infection induced greater levels of both IFN-I and IFN-III transcript levels, and of ISGs such as ISG15 and CXCL-10 (Fig. 4B). However, the increase in IFN/ISG transcript levels, ∼2-3 fold, is somewhat reduced compared to those prior results with SARS-CoV-1 and MHV-JHM Mac1 mutant viruses (26, 32). This differences in IFN induction between WT and Mac1 deleted/mutant viruses between different CoVs could be due to alterations in the functions or abundance of other CoV-encoded IFN repressing proteins expressed by SARS-CoV-2.

**Fig. 4.**
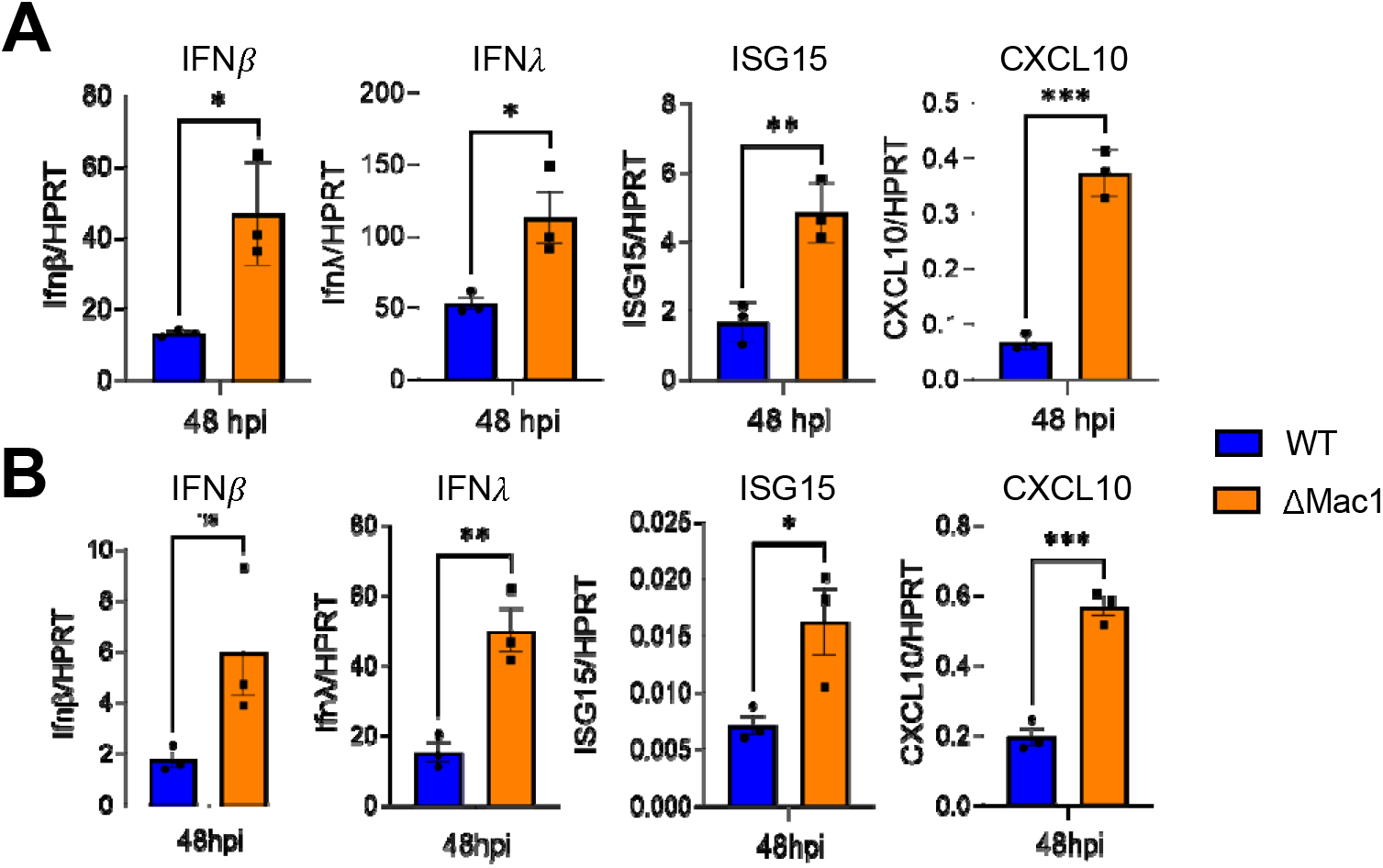
ΔMac1 induces increased IFN and cytokines responses compared to WT SARS-CoV-2 in cell culture. Calu3 (A) and A549-ACE2 (B) cells were infected with SARS-CoV-2 WT and ΔMac1 at an MOI of 0.1 PFU/cell and total RNA was collected 48 hpi. IFN-β, IFN-λ, ISG15 and CXCL10 levels were determined by qPCR using ΔCt method with primers listed in table S2 and normalized to HPRT mRNA levels. The data show one experimental representative of three independent experiments with *n* = 3 for each experiment.

### SARS-CoV-2 ΔMac1 is highly attenuated in K18-ACE2 mice

We next tested the ability of SARS-CoV-2 WT and ΔMac1 to cause disease in K18-ACE2 C57BL/6 mice, a lethal animal model of SARS-CoV-2 infection. Following intranasally inoculation of 2.5×10^4^ PFU WT SARS-CoV-2, we observed 100% morbidity and mortality. In contrast, SARS-CoV-2 ΔMac1 infection did not cause any weight loss or lethality, indicating extreme attenuation (Fig. 5A-B). When analyzing the infected lungs by hematoxylin and eosin staining, we noted significantly higher levels of bronchointerstitial pneumonia, inflammation, and edema and fibrin in WT SARS-CoV-2 infected lungs compared to ΔMac1 virus infected lungs (Fig. 5C). We then compared the WT and ΔMac1 SARS-CoV-2 loads in infected lungs and found that by day 1 post-infection there was a significant reduction in viral titers (Fig. 5E) and viral genomic RNA (gRNA) (Fig. 5F) of ∼ 1 log in the lungs of ΔMac1 infected mice compared to WT SARS-Cov-2 infected lungs. The difference in viral load between WT and ΔMac1 increased to 2.5 logs by day 3, and by day 7 ΔMac1 was effectively cleared from the lungs while WT virus was still present at about 10^5^ PFU in the lung (Fig. 5E-F). In contrast, viral loads in the brain were very low until after day 3 post-infection, though WT virus was present at low levels in most mice by day 7, whereas ΔMac1 titers were below the detection limit at all days tested. (Fig. S4A). Further, there was no significant difference in brain pathology between WT and ΔMac1 infected mice (Fig. S4B), indicating that brain infection and pathology did not significantly contribute to the weight loss and mortality of WT virus infected mice.

**Fig. 5.**
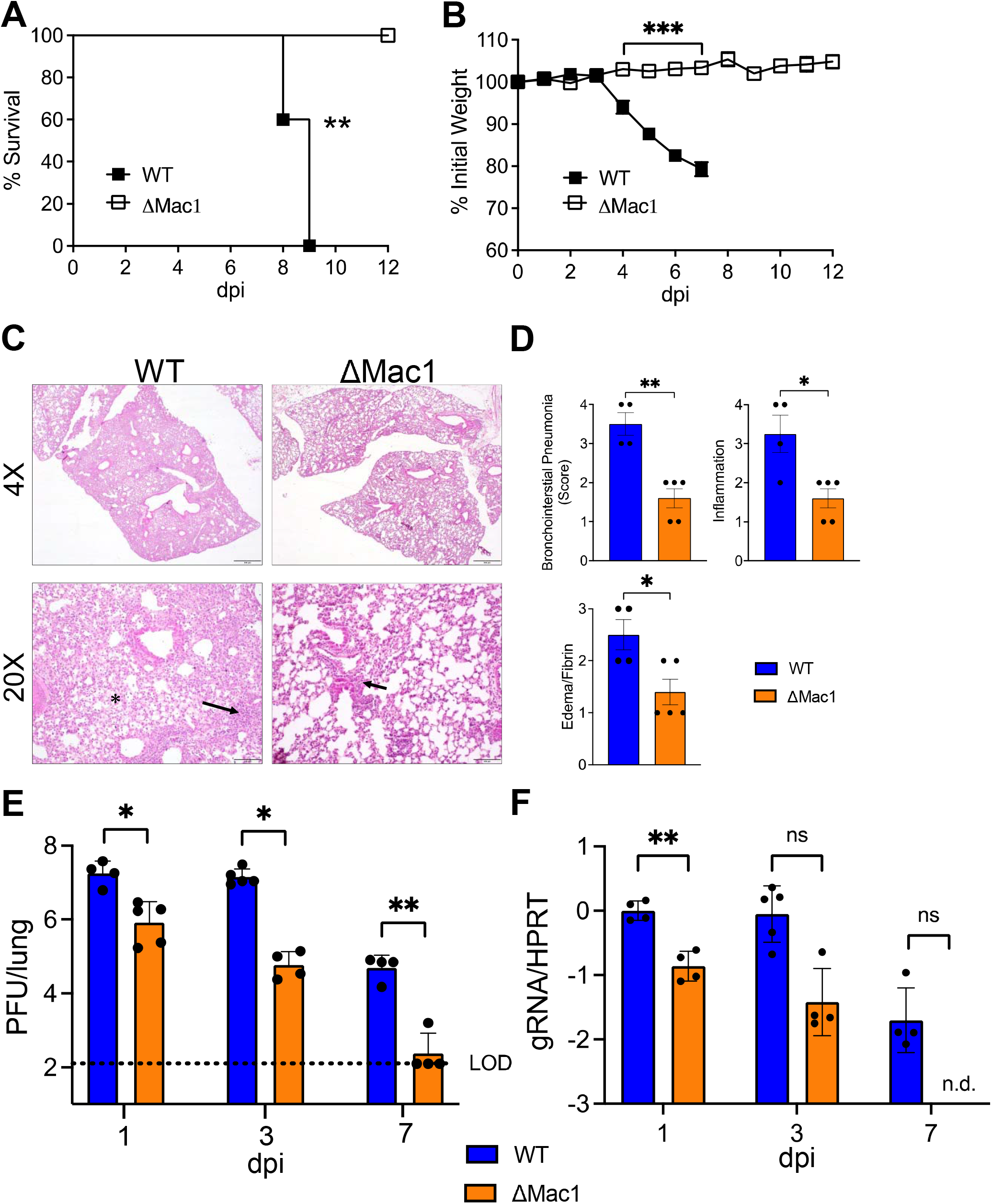
ΔMac1 is highly attenuated in K18-ACE2 mice. (A-B) K18-ACE2 C57BL/6 mice were infected with 2.5×104 PFU of WT or ΔMac1 SARS-CoV-2 and survival and weight loss were measured over 12 days. (C) Photomicrographs (hematoxylin and eosin stain) of lungs from WT and ΔMac1 infected mice at 7 dpi demonstrating bronchointerstitial pneumonia (black arrow) and edema and fibrin (black asterisk) (D) Mice were scored for bronchointerstitial pneumonia, inflammation, and edema/fibrin deposition. WT n=4; ΔMac1 n=5. (E-F) K18-ACE2 C57BL/6 mice were infected as described above and lung titers (E) and gRNA levels (F) were determined by plaque assay and RT-qPCR with primers specific for nsp12 and normalized to HPRT, respectively. Results are from one experiment representative of two independent experiments. *n* = 4-10 mice per group.

### SARS-CoV-2 ΔMac1 induces a robust innate immune response in the lungs of K18-ACE2 mice

The rapid clearance of ΔMac1 in the lungs of infected mice and prior results with Mac1 SARS-CoV mutant viruses (18) suggested that ΔMac1 would induce a strong innate immune response in mice. To test this possibility, we measured the transcripts of a small panel of IFN and ISGs for their expression following infection of WT and ΔMac1 at 1 day post infection (Fig. 6A). IFN-/*β* and IFN-λ were upregulated by more than 10-fold in ΔMac1 infected lungs, while IFN-γ was not detected. We also observed a ∼2-3-fold increase in several ISGs, such as OAS, ISG15, CXCL10, IL-6, PARP12, and PARP14. These results suggest that the attenuation of ΔMac1 virus could, at least in part, be due to a robust IFN response at the early stages of infection. To get a global view of all the transcriptional changes occurring in the absence of Mac1, we performed RNAseq of whole lung samples collected at day 1 post-infection.

**Fig. 6.**
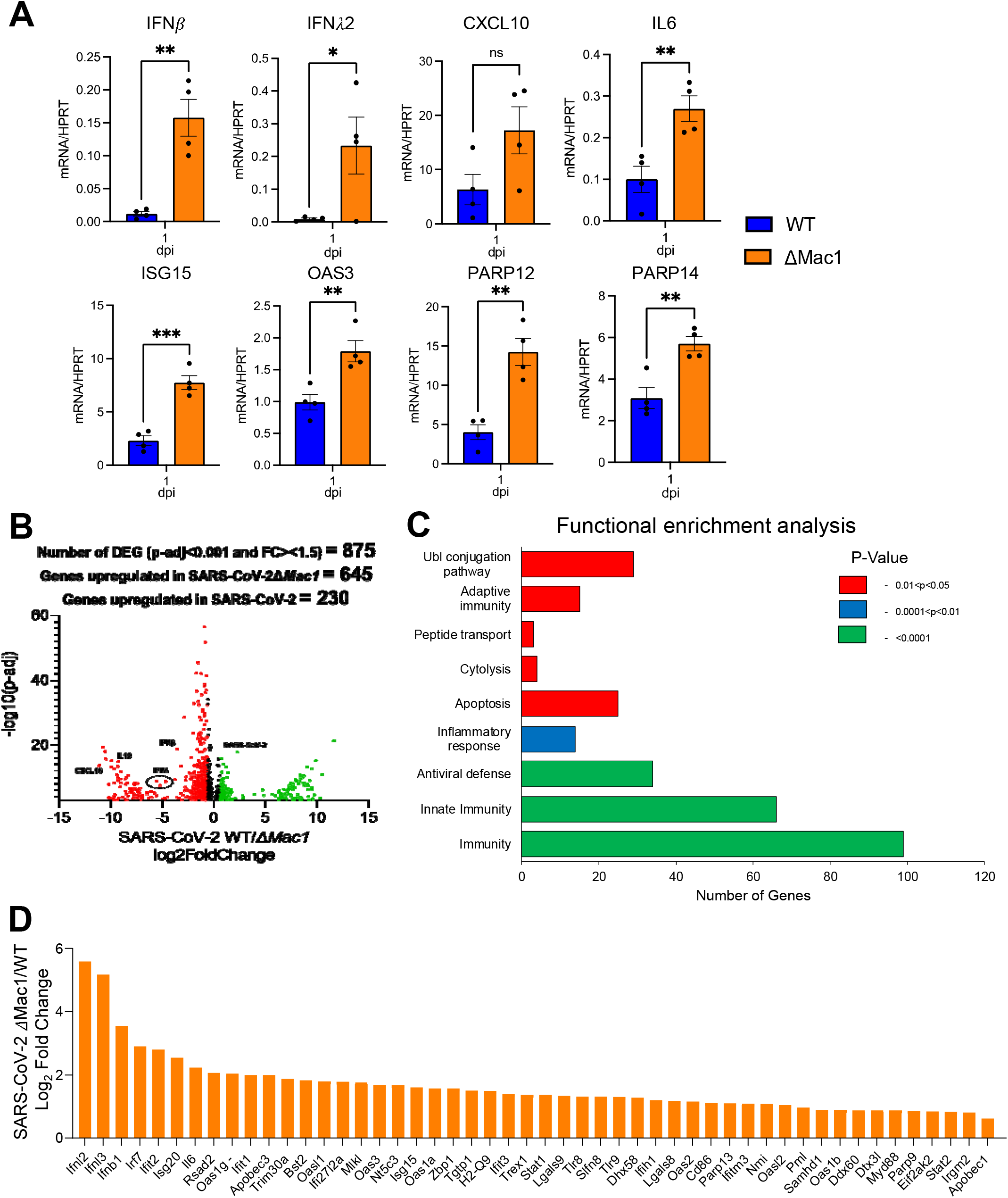
ΔMac1 virus induces a robust innate immune response in the lungs following infection. (A) K18-ACE2 C57BL/6 mice were infected with 2.5×104 PFU of indicated viruses and lungs were harvested at 1 dpi and total RNA was isolated. The relative levels of indicated transcripts were determined by qPCR using the ΔCt method with primers listed in table S2 normalized to HPRT mRNA levels. The results are from one experiment representative of two independent experiments with an *n* = 4-8 mice per group. (B-D) The total RNA from the samples in (A) were analyzed by RNAseq to determine the full transcriptome in the lung following infection. (B) Volcano plot indicating *<u>d</u>*ifferentially *<u>e</u>*xpressed *<u>g</u>*enes (DEGs) between WT and ΔMac1 infected mice. (C) Functional enrichment analysis of biological processes enriched in the transcriptome of in mice infected with ΔMac1 performed using DAVID functional annotation tool. (D) Log_2_ fold change values of genes involved in innate immune response upregulated in mice infected with ΔMac1 compared to WT virus.

Differentially expressed genes were define as having at least 1.5-fold increased expression in either WT or ΔMac1 infected lungs with an adjusted p value of <0.05. In total, we found 645 genes were increased following infection with ΔMac1, and another 230 were increased following WT infection, including viral gRNA, for a total of 875 differentially regulated genes (Fig. 6B).

We then performed a gene ontology analysis, and found that genes related to immunity, innate immunity, and antiviral defense were the pathways that were most significantly upregulated in ΔMac1 infected lungs (Fig. 6C). In addition, genes in the categories of adaptive immunity, ubiquitin conjugation, inflammatory responses, peptide transport, cytolysis, and apoptosis were also significantly upregulated in ΔMac1 infected lungs (Fig. 6C). We then looked at the individual expression of a panel of ISGs and found that most ISGs were increased between 2 and 4-fold in ΔMac1 when compared to WT virus infection, while IFN-/*β* and IFN-λ were increased more than 10-fold (Fig. 6D, Fig. S5). In total, we have found that Mac1 is required for SARS-CoV-2 to block the innate immune response during SARS-CoV-2 infection in mice.

### SARS-CoV-2 ΔMac1 infection results in reduced myeloid cell accumulation in the lungs

Next, we assessed the impact of WT and ΔMac1 virus infection on the recruitment of innate immune cells, specifically inflammatory monocytes and neutrophils, into the lung that might contribute differential lung inflammation and disease severity. Inflammatory monocytes were found to contribute to disease severity in SARS-CoV-1 and MERS-CoV infected mice by promoting the production of TNFa and increased T cell apoptosis (44, 45). Previously, IFN-I was shown to enhance inflammatory monocyte accumulation in the lung, though this was due to IFN-I production in the later stages of SARS-CoV-1 replication (44). However, earlier exogenous addition of IFN-I reduced inflammatory monocyte infiltration following MERS-CoV infection was shown to reduce the number of inflammatory monocytes (45). Thus, we hypothesized that the early IFN-I and IFN-III induction by ΔMac1 would result in fewer infiltrating inflammatory immune cells. Following infection with ΔMac1, we observed a substantial reduction in both the percentage and total number of inflammatory monocytes at both 3 and 7 days after infection (Fig. 7A), which could also play a role in the attenuation of the disease severity. Neutrophils were slightly increased in percentage in ΔMac1 infected lungs at day 3 but had similar total numbers when compared to WT virus infection (Fig. 7B). However, by day 7 there was a significant reduction in the total number of neutrophils in ΔMac1 infected lungs (Fig. 7B).

**Fig. 7.**
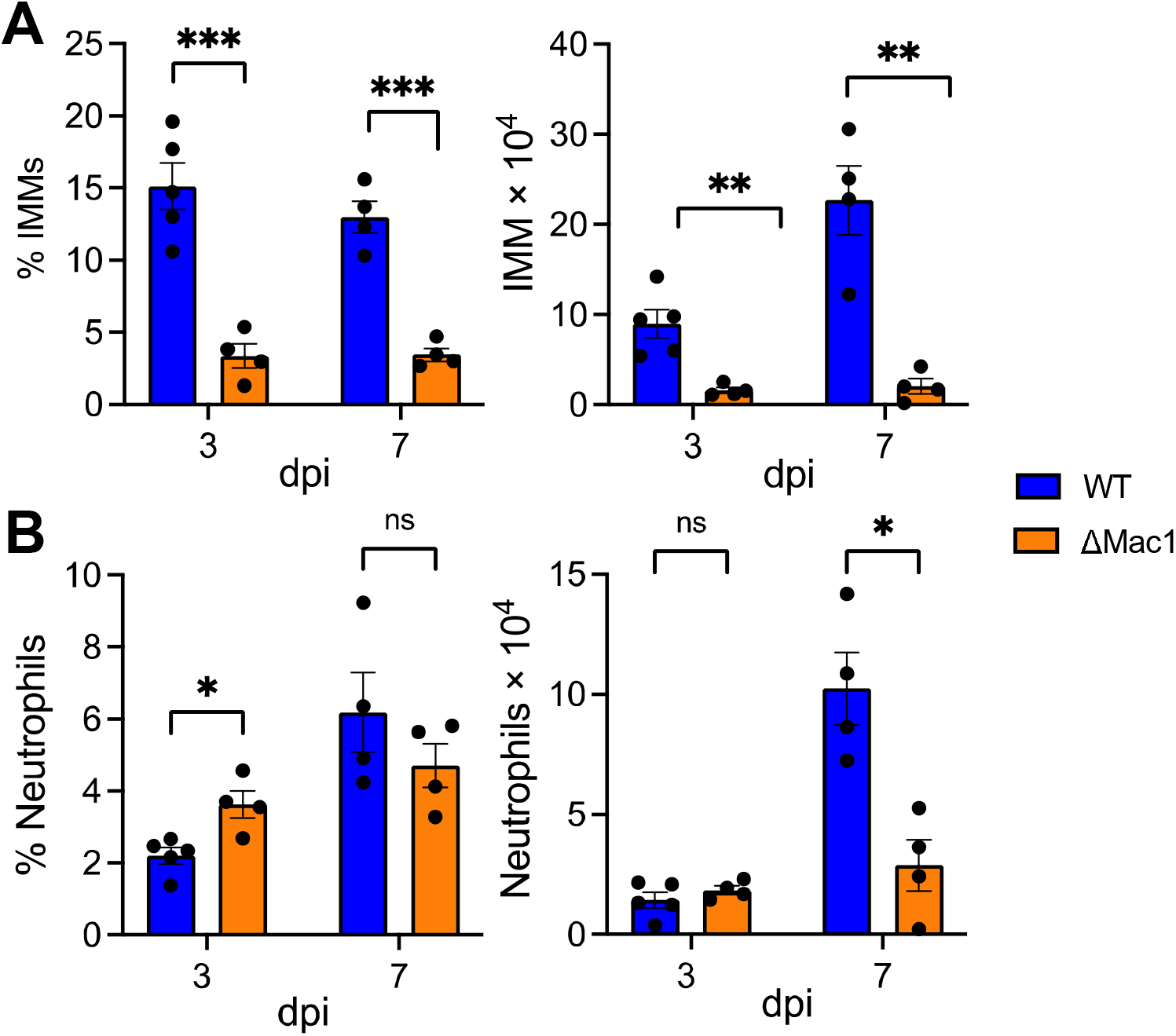
ΔMac1 virus infection results in reduced inflammatory monocytes and neutrophils. (A-B) K18-ACE2 C57BL/6 mice were infected as described above and lungs were harvested at the indicated days post-infection, and the percentages and total numbers of infiltrating inflammatory monocytes (A) and neutrophils (B) were determined by flow cytometry. Data are derived from the results of 1 experiment representative of 2 independent experiments performed with 4-5 mice/group/experiment.

Overall, our results indicate that the absence of Mac1 promotes a strong IFN response with a reduction in inflammatory cell types that may both play a role in reducing viral loads and preventing disease following infection.

## DISCUSSION

The COVID-19 pandemic caused by SARS-CoV-2, has fueled a new interest in the development of inhibitors targeting viral gene products that block viral replication and pathogenesis, to be used therapeutically to treat patients with COVID-19. This includes the $577 million-dollar Antiviral Drug Development Award (AVIDD) initiated by the NIH in 2021. Thus far, antivirals targeting the viral polymerase (nsp12) and protease (nsp5) have been approved for clinical use (4-6), however a much larger anti-CoV drug portfolio is clearly needed to target SARS-CoV-2 and effectively respond to novel CoV outbreaks in the future. One of the CoV-encoded proteins that has received increased attention as a potential drug target is the conserved macrodomain, now called Mac1 (46). Multiple groups have initiated drug development programs targeting Mac1, all utilizing biochemical assays that can be used to screen for compounds that inhibit either Mac1-ADP-ribose binding or Mac1 ADP-ribosylhydrolase activity (34-41). Currently, the top Mac1 inhibitors identified to date have IC_50_ values in these biochemical assays ranging from ∼0.5-10 µM. However, none of these compounds have been tested for their ability to inhibit virus replication or pathogenesis in cell culture or in mice. While the current body of literature indicates that Mac1 is important for the replication and pathogenesis of MHV and SARS-CoV-1 in mice (18, 26, 31), no study has yet evaluated how Mac1 impacts the replication of SARS-CoV-2, which is critical for the ability to interpret inhibitor studies.

Prior results in our lab had indicated that Mac1 is critical for the replication of MHV-JHM, as at least two Mac1 mutant recombinant BACs failed to produce infectious virus (33). However, one of these mutations, G1439V, did replicate after acquiring a second site mutation in the residue immediately preceding it, A1438T. To confirm these results, we created a complete deletion of Mac1 in the MHV-JHM BAC and again found that we could not recover infectious virus (Fig.S1A-B). We hypothesized that perhaps a Mac1 deletion may be detrimental across CoVs, so we then created MERS-CoV and SARS-CoV-2 ΔMac1 recombinant BACs. Surprisingly, we were unable to recover infectious virus from the MERS-CoV ΔMac1 BAC but easily recovered infectious SARS-CoV-2 ΔMac1 (Fig. S1A-B). While we haven’t tested a Mac1 deletion in SARS-CoV-1, the near WT replication of the same G-V mutation in MHV described above indicates that Mac1 is likely non-essential for SARS-CoV-1 as well (18). This near absolute requirement for Mac1 in some CoV species but not in others was surprising, but not without precedent. For instance, a recombinant virus with an E protein deletion was viable with only a mild replication defect in SARS-CoV-1 (47), but an E protein deletion in MERS-CoV was unrecoverable unless the virus was propagated on E protein expressing cells (48). Nsp14 ExoN mutations are lethal in MERS-CoV and SARS-CoV-2 but are viable in MHV and SARS-CoV-1 (49). Finally, MHV nsp15 mutant viruses grow very poorly in IFN-competent macrophages (50, 51), while similar mutations in MERS-CoV replicate normally in IFN-competent cells (52). In the cases of E protein and nsp15 the viruses that replicate normally in the absence of these proteins have additional accessory proteins that have overlapping or redundant functions. For instance, the 4a and 4b MERS-CoV proteins were found to have redundant functions with nsp15 in blocking the innate immune response to infection (52). We hypothesize that the Sarbecoviruses may have evolved unique accessory proteins or other domains in the non-structural proteins that have redundant functions with Mac1 in promoting viral replication.

Efforts to identify these proteins with redundant function are ongoing. Regardless, we and others have found that Mac1 is critical for CoVs to replicate efficiently and cause disease in all animal models of infection that have been tested (Fig. 5) (18, 26, 31).

Here we found that there was no defect in the replication of ΔMac1 in Vero E6 and A549 cells and only a modest defect in Calu-3 cells. Given these results, along with modularity of the various domains of nsp3, it is highly unlikely that the complete deletion of Mac1 had a significant effect on the overall structure of nsp3. Despite the lack of a large replication defect of ΔMac1 under normal growth conditions, we found that ΔMac1 had a >1 log defect in IFN-γ, but not IFN-/*β* treated Calu-3 cells (Fig. 5). IFN-γ induces a small number of ISGs compared to IFN-/*β* (43), and we hypothesized that while the PARP enzymes that inhibit Mac1 mutant MHV are upregulated by both types of IFN, other more potent anti-SARS-CoV-2 ISGs are only upregulated by IFN-/*β*, or at least upregulated to a much higher level by IFN-/*β*. We hypothesized that these ISGs might limit viral entry, mitigating the effect of PARP enzymes to specifically target the Mac1 mutant virus during later stages of the viral lifecycle. It will be of interest to identify the specific PARP or PARPs that inhibit ΔMac1 following IFN-γ treatment and the ADP-ribosylated target(s) that contribute to this inhibition. The ability to specifically reduce ΔMac1 replication with IFN-γ could have important implications for Mac1 inhibitor testing.

Replication assays could be developed to assess the ability of the Mac1 inhibitors to reduce virus replication in the absence and presence of IFN-γ to show that the inhibitors are indeed specifically targeting Mac1. It may also be of interest to test the replication of ΔMac1 under conditions of cell stress, such as ER stress or following activation of stress granules, as PARP activity is known to be increased under stress conditions (53, 54).

Similar to SARS-CoV-1, we found that SARS-CoV-2 ΔMac1 induces a robust innate immune response both in cell culture and in mice (Figs. 4,6), further confirming that Mac1 is one of the many potent IFN repressing proteins expressed by CoVs. This innate immune response occurred within one day of infection, and likely before peak replication of the virus. Whole lung RNAseq data identified over 100 genes involved in immunity to virus infection, demonstrating the breadth of the immune response that is triggered following ΔMac1 infection (Fig. 6B-D). We hypothesize that this response at least partially protects mice from disease. We have previously shown that the Mac1 mutant virus, N1347A, causes increased lethality and weight loss in IFNAR knockout mice, and that providing exogenous IFN prior to peak viral replication is highly protective in multiple animal models of CoV infection and protection is mediated by a combination of increasing the abundance of antiviral ISGs, reducing the number of pro-inflammatory monocytes, and increasing adaptive immunity (32, 44, 45). COVID-19 patients with mutations in IFN related genes or that produce antibodies that target IFN have worse outcomes than the general population, demonstrating the importance of IFN in protection from severe COVID-19 (55-57). However, it remains unclear how an early IFN response from a virus contributes to the protection of mice, and potentially humans, from disease. Some important questions include: How is the IFN induced? Is it through MDA5, RIG-I, or other sensors? Which cell type is the major IFN source from, epithelial or plasmacytoid dendritic cells or myeloid cells? If it’s coming from epithelial cells, which type of epithelial cell; nasal, bronchial, or alveolar? Also, are both IFN-I and IFN-III important for protection, or is one of them sufficient? Finally, how do these IFNs shape the overall innate and adaptive immune response? The answers to these and other questions could have important implications for developing vaccines or therapeutics that might stimulate better and longer lasting immunity and reduce incidence of SARS-CoV-2 spread and disease in vulnerable populations.

## METHODS

### Cell culture and reagents

Vero E6, Huh-7, Baby Hamster Kidney cells expressing the mouse virus receptor CEACAM1 (BHK-MVR) (gifts from Stanley Perlman, University of Iowa), and A549-ACE2 cells (a gift from Susan Weiss, University of Pennsylvania), were grown in Dulbecco’s modified Eagle medium (DMEM) supplemented with 10% fetal bovine serum (FBS). Calu-3 cells (ATCC) were grown in MEM supplemented with 20% FBS. Human IFN-/*β* and IFN-γ were purchased from R&D Systems. Cells were transfected with either Polyjet (Amgen) or Lipofectamine 3000 (Fisher Scientific) per the instructions of the manufacturers

### Mice

Pathogen-free K18-ACE2 C57BL/6 mice were purchased from Jackson Laboratories. Mice were bred and maintained in the animal resources facility at the Oklahoma State University. Animal studies were approved by the University of Oklahoma State Institutional Animal Care and Use Committee (IACUC) and met stipulations of the *Guide for the Care and Use of Laboratory Animals*.

### Generation of recombinant pBAC-SARS-CoV-2, pBAC-MERS-CoV, and pBAC-JHMV constructs

All recombinant pBAC constructs were created using Red recombination (58) with several previously described CoV bacterial artificial chromosomes (BACs). These include the WT-SARS-CoV-2 BAC based off the Wuhan-Hu-1 isolate provided by Sonia Zuñiga, Li Wang, Isabel Sola and Luis Enjuanes (CNB-CSIC, Madrid, Spain) (59), a MERS-CoV BAC based of the EMC isolate (48), and an MHV BAC based on the JHMV isolate (26). All constructs were engineered using a Kan^r^-I-SceI marker cassette for dual positive and negative selection as previously described (see primers in Table S1) (60). Both forward and reverse primers were designed to include a 40bp region upstream of Mac1 to facilitate the deletion of Mac1 by recombination (Table S1). Final BAC DNA constructs were confirmed by restriction enzyme digestion, PCR, and direct sequencing for the identification of correct clones.

### Reconstitution of recombinant pBAC-SARS-CoV-2-derived virus

All work with SARS-CoV-2 and MERS-CoV was conducted in either the University of Kansas or the Oklahoma State University EHS-approved BSL-3 facilities. To generate SARS-CoV-2 or MERS-CoV viruses, approximately 5×10^5^ Huh-7 cells were transfected with 2 *µ*g of purified BAC DNA using Lipofectamine 3000 (Fisher Scientific) as a transfection reagent. SARS-CoV-2 generated from these transfections (p0) was then passaged in Vero E6 cells to great viral stocks (p1). All p1 stocks were again sequenced to confirm that they retained the correct Mac1 deletion and to ensure the furin cleavage site had not been lost (for primers see Table S2). To generate MHV-JHM and MERS-CoV virus, approximately 5×10^5^ BHK-MVR cells were transfected with 1 *µ*g of purified BAC DNA using PolyJet™ Transfection Reagent (SignaGen). In the case of MHV-JHM, an additional 1 *µ*g of N protein-expressing plasmid was co-transfected with genomic BAC DNA.

### Virus infection

Vero-E6, A549-ACE2, or Calu-3 cells were infected at the indicated MOIs. For Calu-3 cells, trypsin-TPCK (1 *µ*g/ml) was added to the medium at the time of infection. All infections included a 1-hour adsorption phase, except for Calu-3 cells where the adsorption phase was increased to 2 hrs. Infected cells and supernatants were collected at indicated time points and titers were determined on Vero E6 cells. For IFN pre-treatment experiments, human IFN-/*β* and IFN-γ were added to Calu-3 cells 18-20 hours prior to infection and were maintained in the culture media throughout the infection. For animal infections, 12-16-week-old K18-ACE2 C57BL/6 female mice were lightly anesthetized using isoflurane and were intranasally infected with 2.5×10^4^ PFU in 50 µl DMEM. To obtain tissue for virus titers, mice were euthanized at different days post challenge, lungs or brains were removed and homogenized in phosphate buffered saline (PBS) and titers were determined on Vero E6 cells.

### Immunoblotting

Total cell extracts were lysed in sample buffer containing SDS, protease and phosphatase inhibitors (Roche), *β*-mercaptoethanol, and a universal nuclease (Fisher Scientific). Proteins were resolved on an SDS polyacrylamide gel, transferred to a polyvinylidene difluoride (PVDF) membrane, hybridized with a primary antibody, reacted with an infrared (IR) dye-conjugated secondary antibody, visualized using a Li-COR Odyssey Imager (Li-COR), and analyzed using Image Studio software. Primary antibodies used for immunoblotting included anti-SARS-CoV-2 N (SinoBiological 40143-R001) and GAPDH (Millipore-Sigma G8795) monoclonal antibodies. Secondary IR antibodies were purchased from Li-COR.

### Confocal Immunofluorescence

Calu-3 cells were cultured with approximately 1.4×10^5^ cells per well in 8-well, removable chamber slides (ibidi 80841) and infected with SARS-CoV-2 at an MOI of 1 PFU/cell. At 24 hpi, monolayers were fixed for 20 minutes with ice cold methanol then 10 minutes with 2% paraformaldehyde in HBSS + 0.01% Sucrose (HBSS/Su). Permeabilization with 0.1% Saponin in HBSS/Su was then performed, followed by overnight blocking at 4°C using 3% goat serum in HBSS/Su + Saponin. Primary antibody incubation was conducted for 3 hours at room temperature (1:2,000 α-N protein, Sino Biological 40143-R001; 1:500 α-nsp3, abcam ab283958) followed by a 1 hour, room temperature secondary antibody incubation (1:200 AlexaFluor 555 Goat α-rabbit, Invitrogen A32732). Nuclear stain with 300nM DAPI was performed at room temperature for 30 minutes followed by mounting in Vectashield Vibrance Mounting Medium (Vector Labs H-1700) and storage at 4°C. Images were acquired using an Olympus FV1000 laser-scanning confocal microscope equipped with Fluoview software. Images were z-projected using maximum intensity.

### Semi-quantitative PCR analysis

BAC DNA or infection-derived cDNA was PCR amplified by primers that bind outside of the Mac1 coding sequence. PCR products were analyzed by gel electrophoresis using a LICOR M imager and bands were quantified using Image Studio software and the relative intensity of each band was determined by adding the overall intensity of both bands together and then diving the intensity of each individual band by the total intensity.

### Real-time quantitative PCR (RT-qPCR) analysis

RNA was isolated from cells using Trizol (Invitrogen). Lungs from K18-ACE2 C57BL/6 mice infected with virus were collected at indicated time points and were homogenized in Trizol (Invitrogen) and RNA was isolated using manufacturer’s instructions. cDNA was prepared using MMLV-reverse transcriptase per the manufacturer’s instructions (Thermo Fisher Scientific). qPCR was performed using PowerUp SYBR green master mix (Applied Biosystems) and primers listed in Table S3. Cycle threshold (CT) values were normalized to hypoxanthine phosphoribosyltransferase (HPRT) levels by using the ΔCt method.

### RNAseq

RNA was isolated from K18-ACE2 mice as described above. Library preparation was performed by the University of Kansas Genome Sequencing core facility, using the NEB Next RNA Library kit (NEB) with indexing. RNA-seq was performed using an Illumina NextSeq2000 high-output system with a paired-end reads of 50 bp each. RNAseq data quality was checked using FastQC analysis pipeline. Samples had a minimum of 16 million reads and a mean quality score (PF) >33. The mouse (C57BL6) transcriptome reference sequence (GCF_000001635.27_GRCm39) and SARS-CoV-2 genome (Accession number - NC_045512.2) were downloaded from NCBI genome collections and appended into a single sequence and used as the reference sequence. RNAseq reads were mapped to the indexed reference sequence using kallisto v0.44.0. Transcripts per kilobase per million mapped reads (TPM) and read counts per transcript were extracted from the kallisto output. TPM values and read counts for all transcripts from each gene were summed to obtain gene-level expression estimates, and the counts per gene were then rounded to the nearest integer. For a given sample, we only considered genes with at least 50 mapped reads total across all replicates from the samples. DESeq2 was used to identify DEGs between the SARS-CoV-2 WT and ΔMac1 infected samples using simply “treatment” as a factor. DEGs were identified based on the false-discovery rate corrected P-value (P_ADJ_) and log_2_-fold-change of (log_2_FC) between the samples. Genes were considered upregulated in a SARS-CoV-2-infected sample if P_ADJ_ < 0.05 and log_2_FC > 0.6, which is nearly equivalent to a 1.5-fold increase. Similarly, genes were considered downregulated if P_ADJ_ < 0.05 and log_2_FC < −0.6, or a 1.5-fold decrease. DEGs were subjected to gene ontology analysis using the Database for Annotation, Visualization and Integrated Discovery (DAVID: https://david.ncifcrf.gov/). Gene lists were analyzed for biological processes that were significantly enriched with PL□<□0.05 and displayed as a clustered bar graph.

### Lung cell preparation and flow cytometry

For phenotypic analyses of lung infiltrating immune cells, lungs collected at different days post-infection, PBS perfused lungs (left lobe were cut into small pieces, treated with collagenase-D and DNAse1 for 30 minutes at room temperature, followed by homogenization of lung pieces using a 3ml syringe plunger flang/thumb rest. Homogenized cells were passed through 70µM strainer to obtain single cell suspension. Isolated single cell suspension was surface immunolabelled for neutrophil (CD45^+^ CD11b^+^ Ly6G^hi^) and inflammatory monocyte (CD45^+^ CD11b^+^ Ly6c^hi^) markers by flow cytometry. For cell surface staining, lung cells were labelled with the following fluorochrome-conjugated monoclonal antibodies: PECy7 α-CD45 (clone: 30-F11); FITC α-Ly6G (clone: 1A8); PE/PerCp-Cy5.5 α-Ly6C (clone: HK1.4); V450 α-CD11b (clone: M1/70); APC α-F4/80 (clone: BM8) (all procured from Biolegend). A detailed cell surface and intracellular immunolabelling protocol for flow cytometry studies are described in our recent publication (61). All fluorochrome-conjugated antibodies were used at a final concentration of 1:200 (antibody: FACS buffer), except for FITC labeled antibodies used at 1:100 concentration.

### Histopathology

The lung lobes were perfused and placed in 10% of formalin. Brain samples were fixed in 10% formalin. The lung lobes and brain were then processed for hematoxylin and eosin (H & E). The lung lesions were blindly scored by an American College of Veterinary Pathology Board-certified pathologist. The lesions were scored on a scale of 0-10% (score 1), 10-40% (score 2), 40-70% (score 3) and >70% (score 4) and cumulative scores were obtained for each mouse. The lesions scored were bronchiointerstitial pneumonia, peribronchial inflammation, edema/fibrin, necrosis, and perivascular inflammation.

### Statistics

A Student’s *t* test was used to analyze differences in mean values between groups. All results are expressed as means ± standard errors of the means (SEM). Differences in survival were calculated using a Kaplan-Meier log-rank test. P values of ≤0.05 were considered statistically significant (*, P≤0.05; **, P≤0.01; ***, P≤0.001; ****, P ≤0.0001; n.s., not significant).

### Data and materials availability

All the RNAseq reads data are deposited in NCBI under the BioProject ID PRJNA928501 and BioSample ID SAMN32942656 and SAMN32942675 and will be made public upon publication or August 31st 2023, whichever comes first.

## ACKNOWLEDGEMENTS

We thank members of the Davido laboratory at KU for valuable discussion, Stanley Perlman and Susan Weiss for reagents, and Brian Ackley for assistance with confocal microscopy.

Bioinformatic consultation was provided by the KU Center for Genomics. Research reported in this publication was made possible in part by the services of the KU Genome Sequencing Core which is supported by the National Institutes of Health under award number P30GM145499.

## Funding

National Institutes of Health (NIH) grant P20GM103648 (RC)

National Institutes of Health (NIH) grant 2P01AI060699 (LE)

National Institutes of Health (NIH) grant P20GM113117 (ARF)

National Institutes of Health (NIH) grant K22AI134993 (ARF)

National Institutes of Health (NIH) grant R35GM138029 (ARF)

National Science Foundation (NSF) grant 2135167 (RLU)

University of Kansas General Research Fund (GRF) and Start-up funds (ARF)

NIH Graduate Training at the Biology-Chemistry Interface grant T32GM132061 (CMK) University of Kansas College of Liberal Arts and Sciences Graduate Research Fellowship (CMK)

Government of Spain (PID2019-107001RB-I00 AEI/FEDER, UE) LE European Commission (H2020-SC1-2019, ISOLDA Project nº 848166-2) LE.

## Author contributions

Conceptualization: YMA, SP, RC, ARF

Methodology: YMA, SP, RG, JJOC, CMK, JJP, RLU, SZ, LE, SM, RC, ARF

Investigation: YMA, SP, RG, JJOC, CMK, JJP, DC, CAM, SM, RC, ARF

Visualization: YMA, SP, RG, JJOC, CMK, JJP, SM, RC, ARF

Supervision: RC, ARF

Writing—original draft: ARF, YA, RC, SP

Writing—review & editing: ARF, RC, SP, RLU, SZ, LE

## Competing interests

The University of Kansas-Lawrence has filed a patent application relating to coronavirus live-attenuated vaccines which lists A.R.F and R.C. as co-inventors.

