## Supplemental Tables for "SARS-CoV-2 Mac1 is required for IFN antagonism and efficient virus replication in mice"

**Supplemental Material**

**Supplemental Tables**

Table S1. Linear Recombination Primers for Engineering Mac1 Deletion BACs

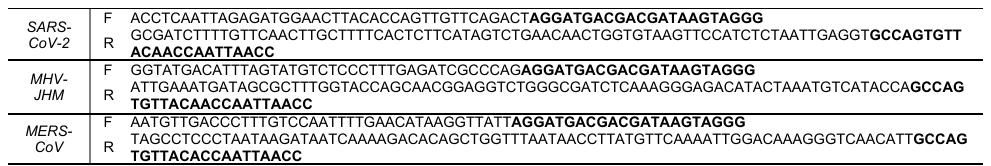

*Letters in Bold indicate Kan^R^-I sequence*

Table S2. Sequencing Primers

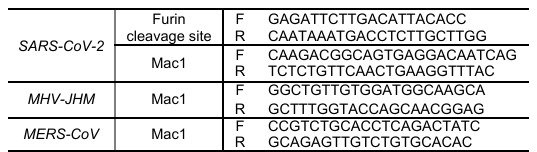

Table S3. qPCR Primers

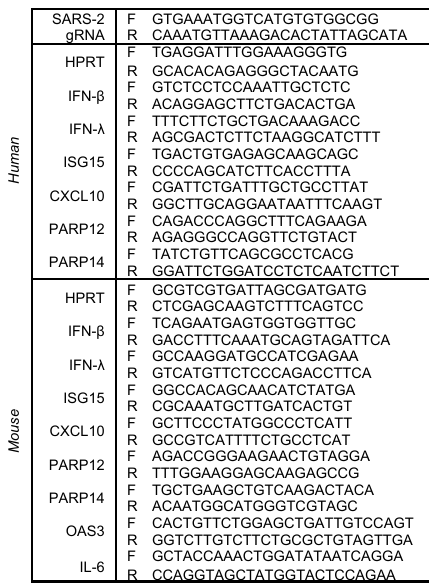
