## Supplemental Figures for "SARS-CoV-2 Mac1 is required for IFN antagonism and efficient virus replication in mice"

### Slide 1
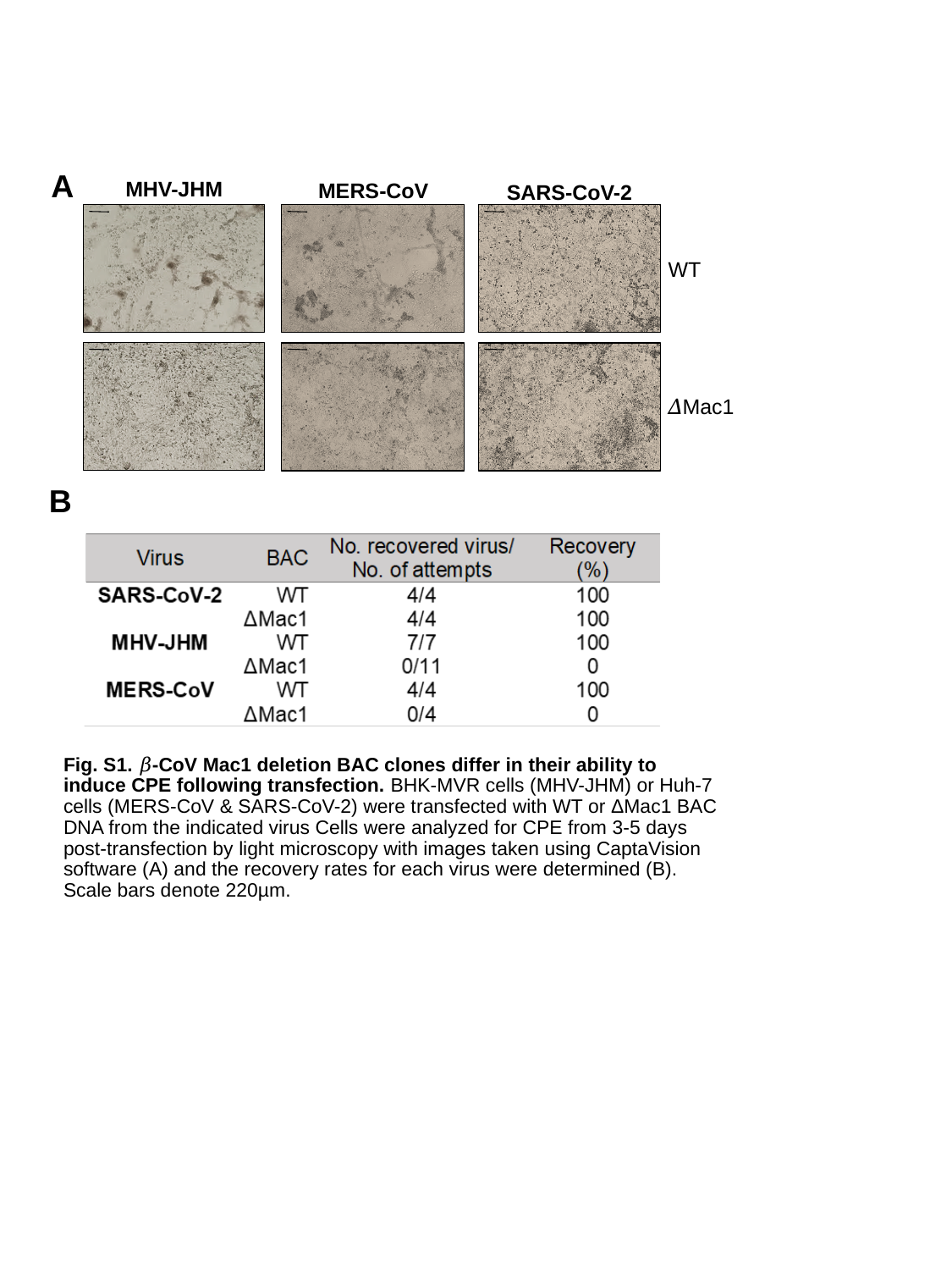

A
MHV-JHM
MERS-CoV
SARS-CoV-2
WT
𝛥Mac1
B
#### Fig. S1. 𝛽-CoV Mac1 deletion BAC clones differ in their ability to induce CPE following transfection. BHK-MVR cells (MHV-JHM) or Huh-7 cells (MERS-CoV & SARS-CoV-2) were transfected with WT or ΔMac1 BAC DNA from the indicated virus Cells were analyzed for CPE from 3-5 days post-transfection by light microscopy with images taken using CaptaVision software (A) and the recovery rates for each virus were determined (B). Scale bars denote 220µm.

### Slide 2
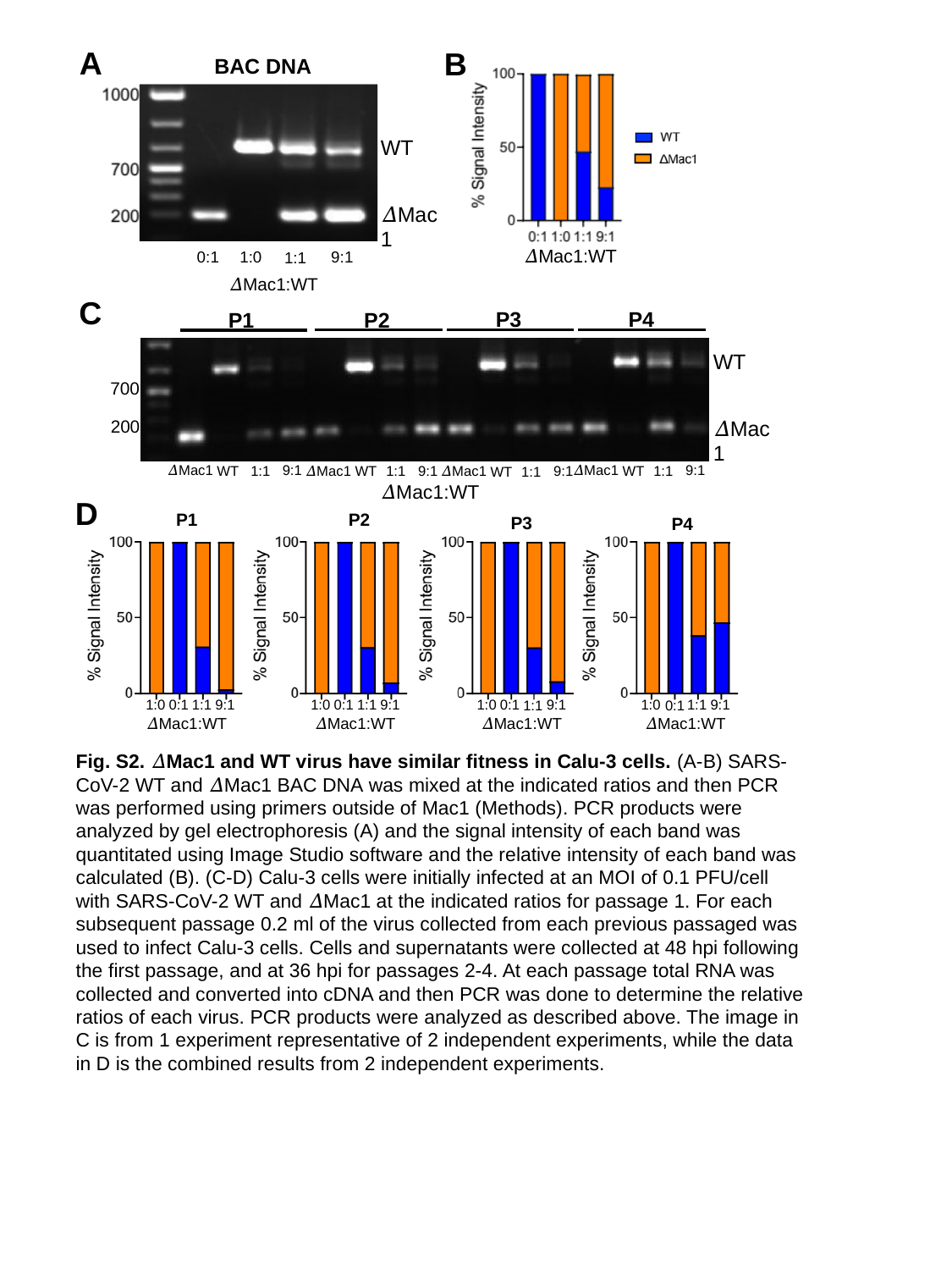

A
B
BAC DNA
WT
𝛥Mac1
𝛥Mac1:WT
0:1
9:1
1:0
1:1
𝛥Mac1:WT
C
P4
P3
P2
P1
WT
700
𝛥Mac1
200
9:1
𝛥Mac1
9:1
𝛥Mac1
WT
𝛥Mac1
9:1
WT
1:1
1:1
WT
1:1
𝛥Mac1
9:1
WT
1:1
𝛥Mac1:WT
D
P1
P2
P3
P4
1:1
1:0
9:1
0:1
1:0
9:1
1:1
0:1
1:0
9:1
1:1
0:1
1:0
9:1
1:1
0:1
𝛥Mac1:WT
𝛥Mac1:WT
𝛥Mac1:WT
𝛥Mac1:WT
Fig. S2. 𝛥Mac1 and WT virus have similar fitness in Calu-3 cells. (A-B) SARS-CoV-2 WT and 𝛥Mac1 BAC DNA was mixed at the indicated ratios and then PCR was performed using primers outside of Mac1 (Methods). PCR products were analyzed by gel electrophoresis (A) and the signal intensity of each band was quantitated using Image Studio software and the relative intensity of each band was calculated (B). (C-D) Calu-3 cells were initially infected at an MOI of 0.1 PFU/cell with SARS-CoV-2 WT and 𝛥Mac1 at the indicated ratios for passage 1. For each subsequent passage 0.2 ml of the virus collected from each previous passaged was used to infect Calu-3 cells. Cells and supernatants were collected at 48 hpi following the first passage, and at 36 hpi for passages 2-4. At each passage total RNA was collected and converted into cDNA and then PCR was done to determine the relative ratios of each virus. PCR products were analyzed as described above. The image in C is from 1 experiment representative of 2 independent experiments, while the data in D is the combined results from 2 independent experiments.

### Slide 3
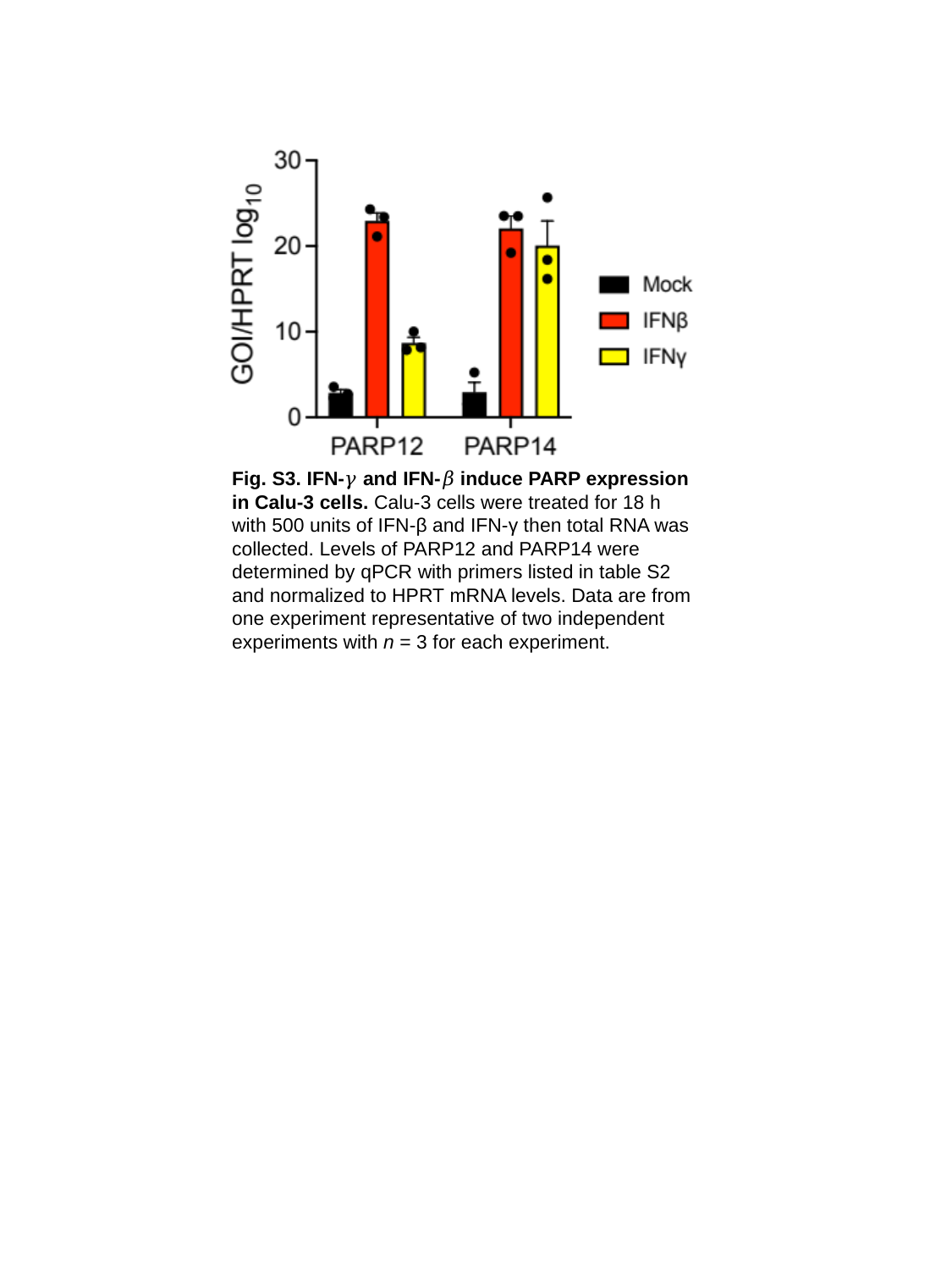

Fig. S3. IFN-𝛾 and IFN-𝛽 induce PARP expression in Calu-3 cells. Calu-3 cells were treated for 18 h with 500 units of IFN-β and IFN-γ then total RNA was collected. Levels of PARP12 and PARP14 were determined by qPCR with primers listed in table S2 and normalized to HPRT mRNA levels. Data are from one experiment representative of two independent experiments with n = 3 for each experiment.

### Slide 4
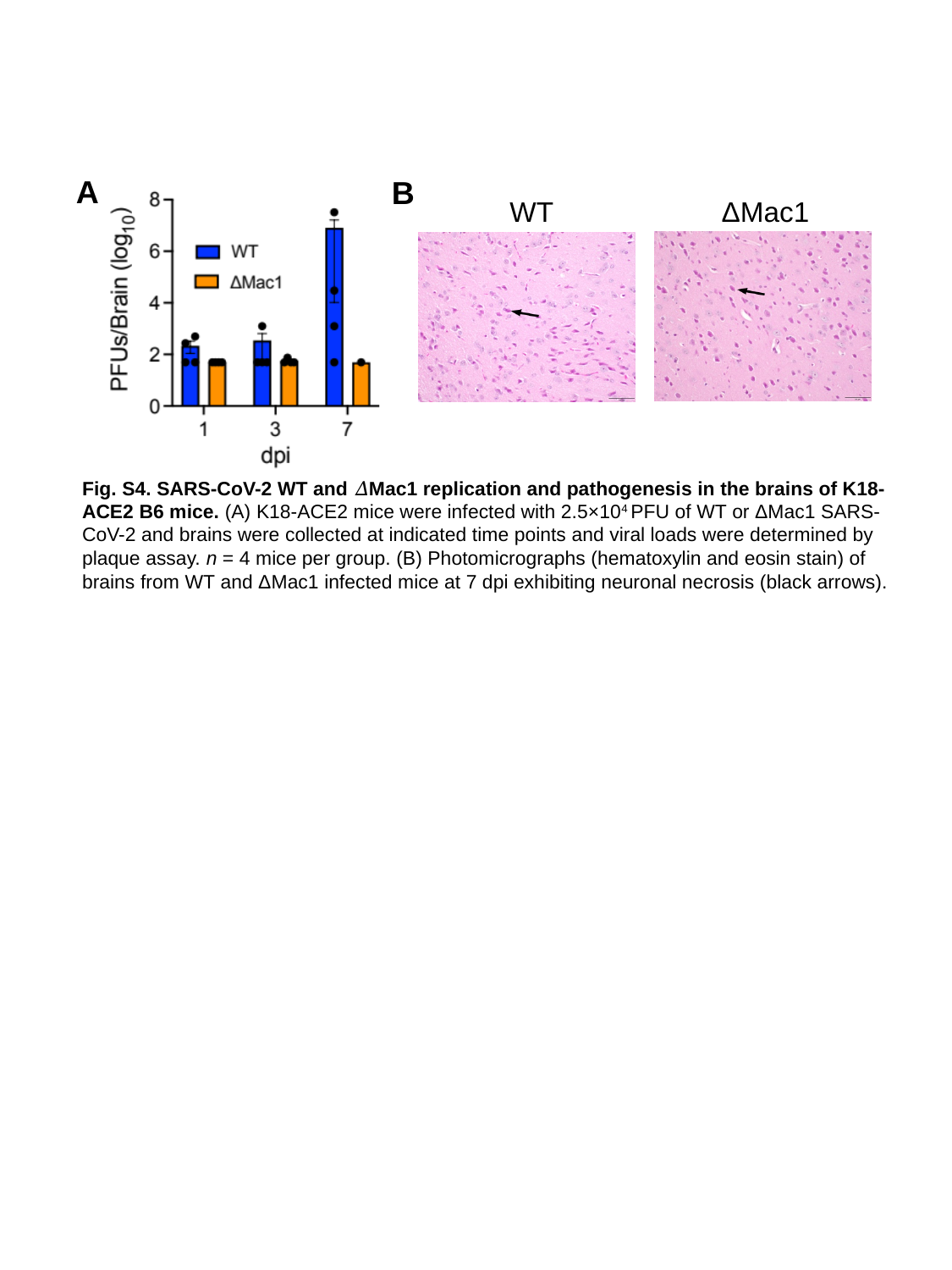

A
B
WT
ΔMac1
#### Fig. S4. SARS-CoV-2 WT and 𝛥Mac1 replication and pathogenesis in the brains of K18-ACE2 B6 mice. (A) K18-ACE2 mice were infected with 2.5×10^4^ PFU of WT or ΔMac1 SARS-CoV-2 and brains were collected at indicated time points and viral loads were determined by plaque assay. n = 4 mice per group. (B) Photomicrographs (hematoxylin and eosin stain) of brains from WT and ΔMac1 infected mice at 7 dpi exhibiting neuronal necrosis (black arrows).

### Slide 5
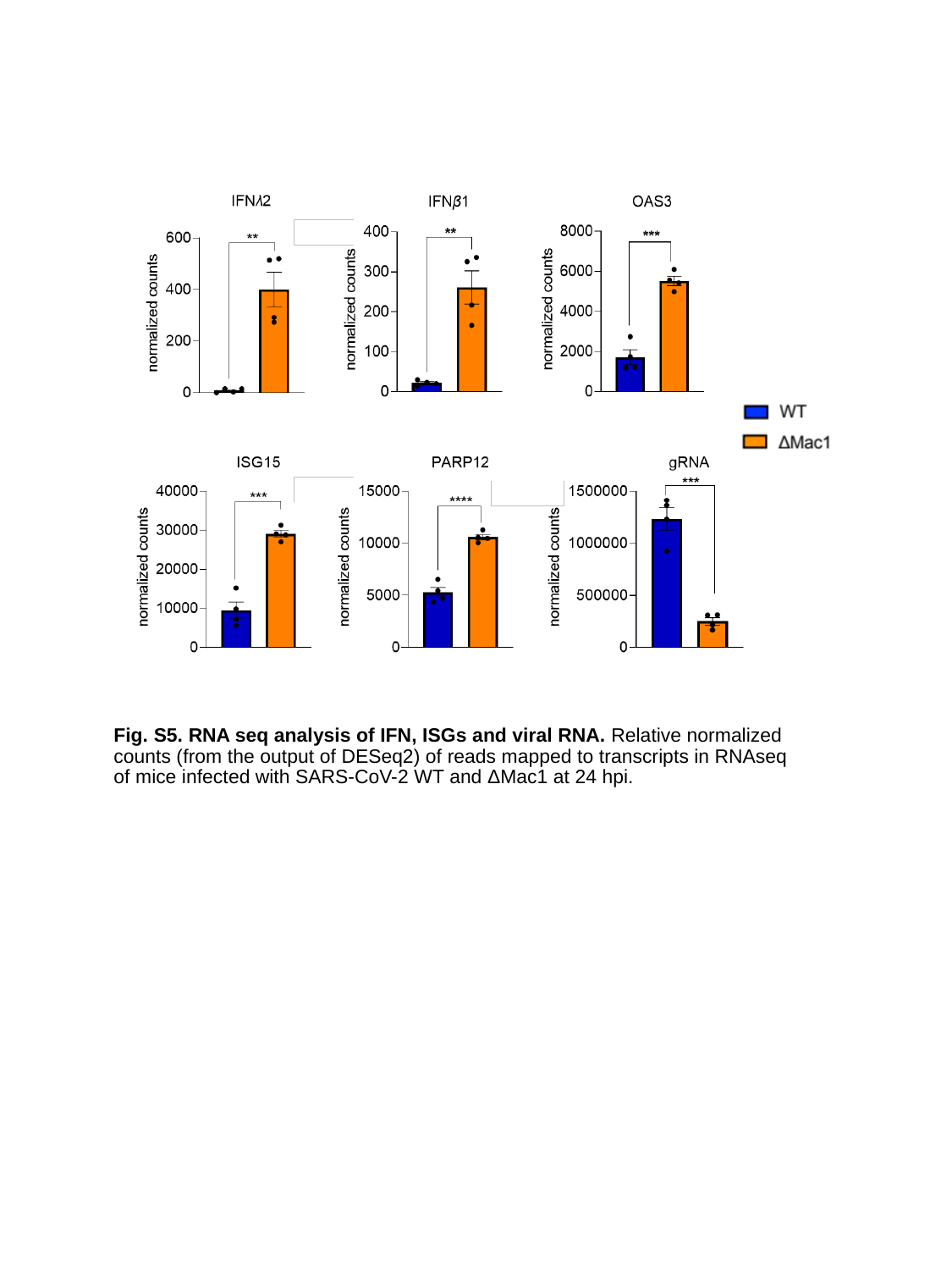

Fig. S5. RNA seq analysis of IFN, ISGs and viral RNA. Relative normalized counts (from the output of DESeq2) of reads mapped to transcripts in RNAseq of mice infected with SARS-CoV-2 WT and ΔMac1 at 24 hpi.
